## supplementary tables for "Single cell RNA sequencing and lineage tracing confirm mesenchyme to epithelial transformation (MET) contributes to repair of the endometrium at menstruation"

**Supplementary Table 1.** Primary and secondary antibodies with associated working dilutions used to detect proteins in mouse uterine tissues

| **Antibody** | **Supplier and Cat. No. (concentration)** | **Dilution in NGS** |
|---|---|---|
| **Primary antibodies** | | |
| Monoclonal rabbit anti-PDGFRβ | Abcam, ab32570 (0.15 mg/ml) | 1/1000 |
| Monoclonal rabbit anti-CD146 | Abcam, ab75769 (0.123mg/ml) | 1/1000 |
| Polyclonal rabbit anti-NG2 | Abcam, ab12905 | 1/600 |
| Polyclonal rabbit anti-CD31 | Abcam, ab28364 (1mg/ml) | 1/500 |
| Polyclonal rabbit anti-EpCAM | Abcam, ab71916 (1mg/ml) | 1/2000 |
| Polyclonal rabbit anti-PDGFRA | Santa Cruz Biotechnology, sc-338 (100µg/ml) | 1/1000 |
| **Secondary antibodies** | | |
| Goat F(ab) anti-rabbit IgG H&L (HRP) | Abcam, ab7171 (1mg/ml) | 1/500 |

**Supplementary Table 2.** Flow cytometry antibodies selected and optimised to interrogate mesenchymal cell populations in murine uterus

| **Antibody** | **Supplier Cat. No. (concentration)** | **Excitation λ (nm)** | **Emission λ (nm)** | **Dilution** |
|---|---|---|---|---|
| BV421 anti-mouse CD31 | BioLegend 102423 (0.2mg/ml) | 405 | 421 | 1/200 |
| BV421 anti-mouse CD45 | Fischer Scientific BDB560501 (0.2mg/ml) | 404 | 448 | 1/100 |
| APC anti-mouse CD146 | BioLegend 134712 (0.2mg/ml) | 650 | 660 | 1/200 |
| BV605 anti-mouse EpCAM | BioLegend 118227 (0.2mg/ml) | 405 | 603 | 1/400 |
| FITC anti-mouse CD90 | Invitrogen 11-0902-82 (0.5 mg/ml) | 490 | 525 | 1/100 |
| DAPI (4',6-Diamidino-2-Phenylindole,Dilactate) | Fischer Scientific D3571 (10mg powder reconstituted to a 0.1µg/µl working solution) | 358 | 461 | 1/1000 |
