## supplementary tables and legens for "Single cell RNA sequencing and lineage tracing confirm mesenchyme to epithelial transformation (MET) contributes to repair of the endometrium at menstruation"

**Supplementary Figures with legends**

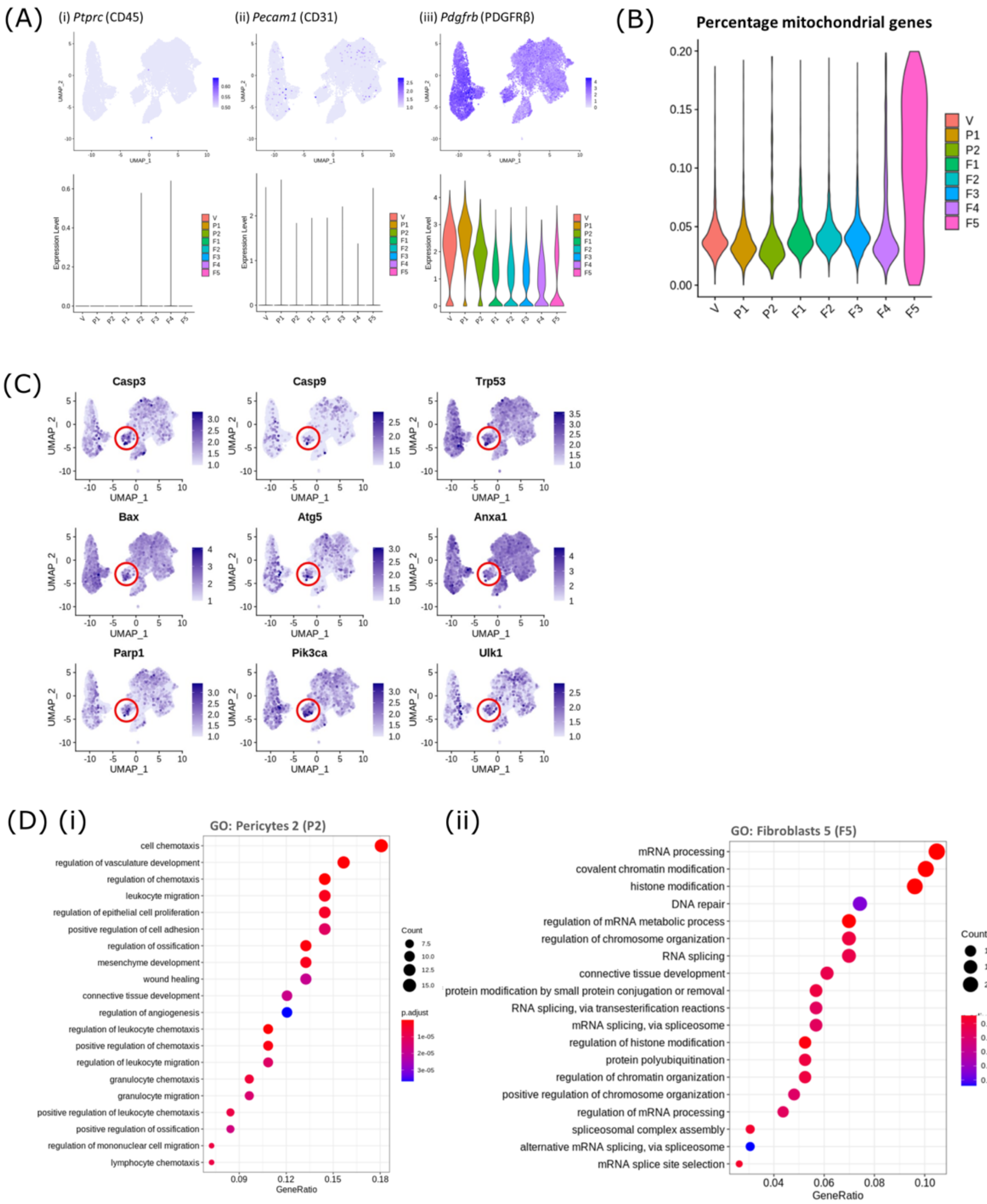


**Supplementary Figure 1. (A)** Gene expression plots: expression of *Pecam1* (CD31), *Ptprc* (CD45) and *Pdgfrb* (PDGFRβ) in the 8 mesenchymal cell clusters confirming that the population of cells isolated for the scRNAseq study had no contamination with endothelial or immune cells. **(B)** Violin plot: percentage of mitochondrial genes expressed by mesenchymal cell clusters indicative of damaged/dying cells; V-vascular smooth muscle cells (vSMCs), P1-pericytes 1, P2-pericytes 2, F1-fibroblasts 1, F2-fibroblasts 2, F3-fibroblasts 3, F4- fibroblasts 4, F5- fibroblasts 5. **(C)** Gene expression plots: expression of apoptosis-associated genes *Casp3*, *Casp9*, *Trp53, Bax, Atg5, Anxa1, Parp1, Pik3ca* and *Ulk1* note high expression in F5. **(D)** Dot plot: clusterProfiler gene ontology (GO) analysis of biological functions associated with the DEgene signature of **(i)** pericytes 2 (P2) and **(ii)** fibroblasts 5 (F5) (dot size: gene ratio, number of genes in data/number of genes associated with GO term; dot colour: p-value representing the enrichment score). Note P2 high for chemotaxis.

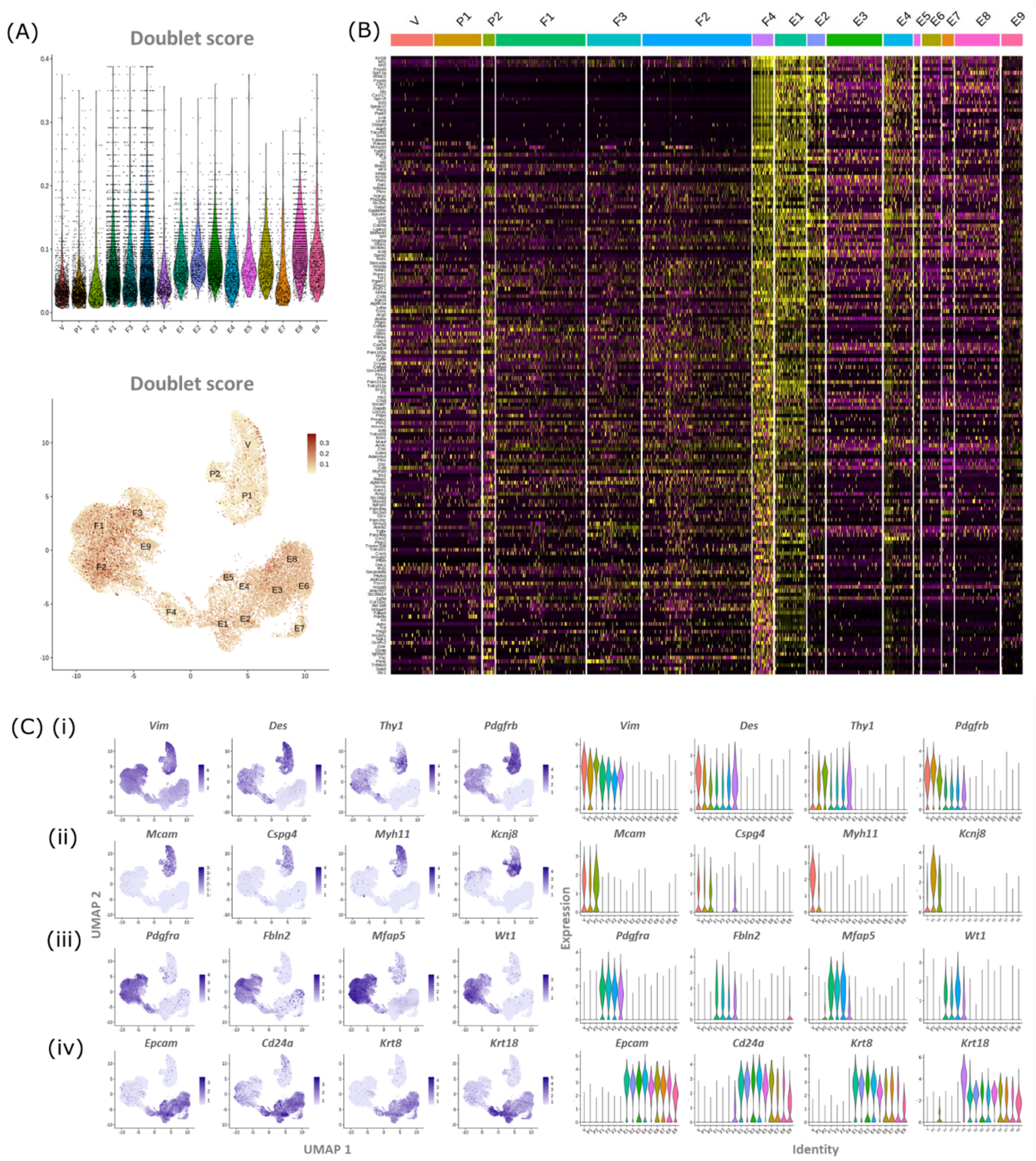


**Supplementary Figure 2. (A)** Expression of canonical markers by GFP+ mesenchymal cells and GFP-EPCAM+ epithelial cells (*Pdgfrb*-BAC-eGFP uterus) for **(i)** mesenchymal (*Vim, Des, Thy1, Pdgfrb*), **(ii)** perivascular (*Mcam, Cspg4, Myh11, Kcnj8*), **(iii)** fibroblast (*Pdgfra, Fbln2, Mfap4, Wt1*), and **(iv)** epithelial (*Epcam, Cd24a, Krt8, Krt18*) cell lineages highlighting the predicted phenotype of the cell clusters. (B) Scaled heatmap (yellow, high; purple, low) displaying differentially expressed genes for cluster F4 when compared to all other clusters (logFC >0.5, pvalue <0.05, Wilcoxon rank-sum test) top is colour coded and named by cluster; V= vascular smooth muscle cells (vSMCs), P1/2= pericytes 1/2, F1-4= fibroblasts 1-4, E1-9= epithelial cells 1-9). Note the transcriptional profile of F4 appears most similar to E1/2.

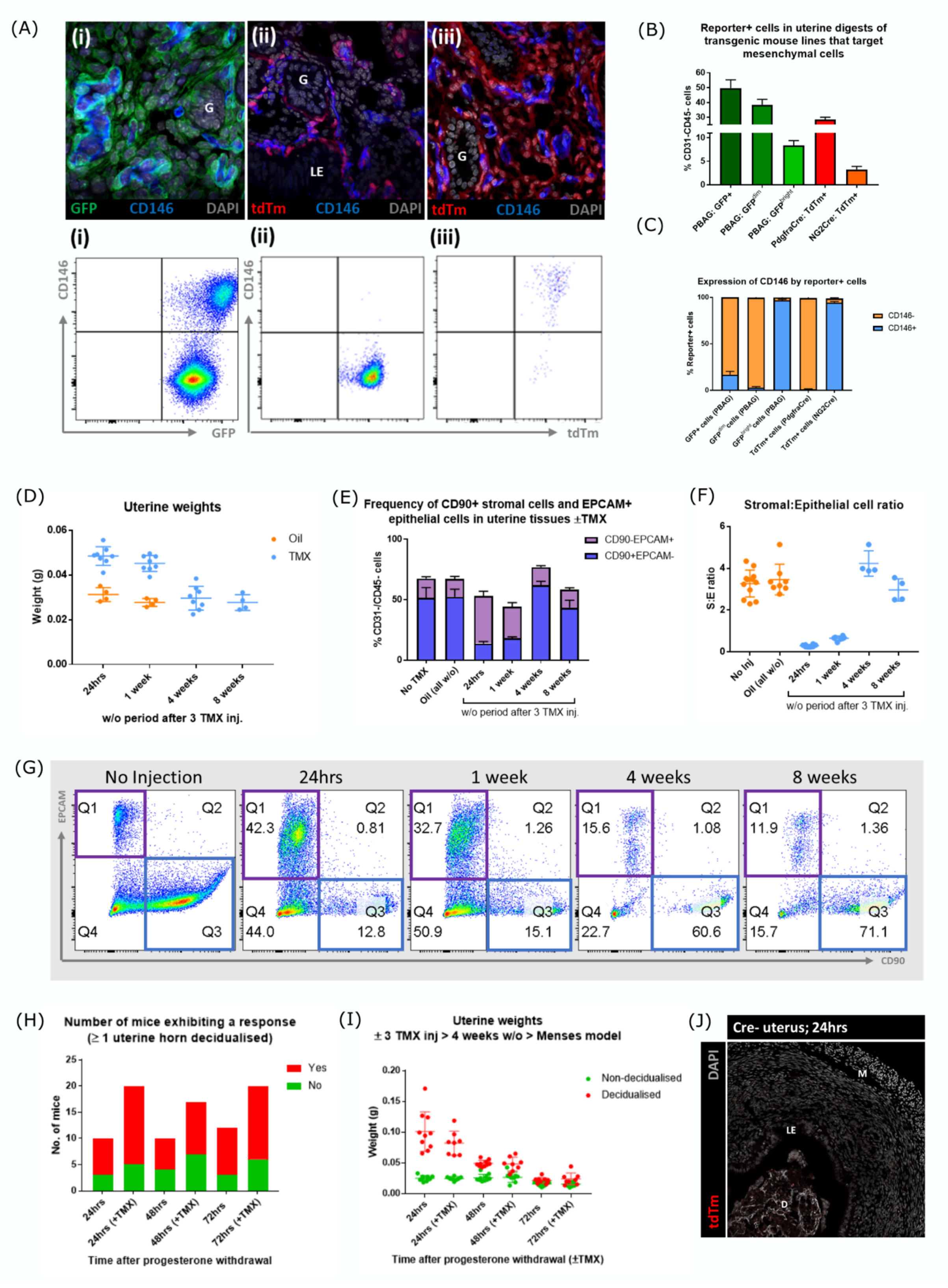


**Supplementary Figure 3. (A)** Expression of canonical pericyte marker CD146 in steady state/control uterine tissues from different transgenic models. **(i)** Two populations of cells are present in *Pdgfrb*-BAC-eGFP uterus: GFP+CD146+ perivascular cells and GFP+CD146- stromal fibroblasts **(ii)** tdTm+ cells in i*Pdgfra*-Cre$^{ERT2}$;*Rosa26-tdTm* uterus are CD146-

(stromal fibroblast phenotype) in contrast **(iii)** tdTm+ cells in iNG2-Cre$^{ERTM}$;*Rosa26-tdTm* uterus are CD146+ (perivascular cell phenotype). **(B)** Bar plot: quantification of flow data analysing the number of reporter+ cells in steady state/control uterine tissues from different transgenic models. **(C)** Bar plot: quantification of flow data analysing the expression CD146 in different transgenic cell populations from control uterine tissues confirming results of immunohistochemistry. **(D)** Bar plot: Uterine horn weights from mice treated with oil or 3x daily TMX analysed after 24h, 1, 4 or 8 weeks. TMX treated animals had uterine weights comparable to the control group by 4 weeks. **(E-F)** Bar plot: quantification of flow data analysing the expression of canonical stromal marker CD90 and epithelial marker EPCAM in uterine tissues from mice treated with oil or TMX analysed after 24h, 1, 4 or 8 weeks. The TMX treated animals had a stromal:epithelial cell ratio comparable to the control groups by 4 weeks. **(G)** Flow cytometry: expression of canonical stromal marker CD90 and epithelial marker EPCAM in uterine tissues from mice treated with oil or TMX – note in line with expectations stimulation of the tissue increased the epithelial compartment (EPCAM+) at 24h and 1 week but this was restored to control levels at 4 weeks. **(H)** Bar plot: Comparison of the number of mice that responded to induction of a decidualisation after initiation of the menstruation model 4 weeks after TMX. **(I)** Bar plot: Uterine horn weights of mice from the mouse model of induced menstruation (non-decidualised and decidualised horns weighed separately) with and without TMX 4 weeks prior to ovariectomy. **(J)** No expression of tdTm detected in the uterus from Cre negative tamoxifen treated mice.

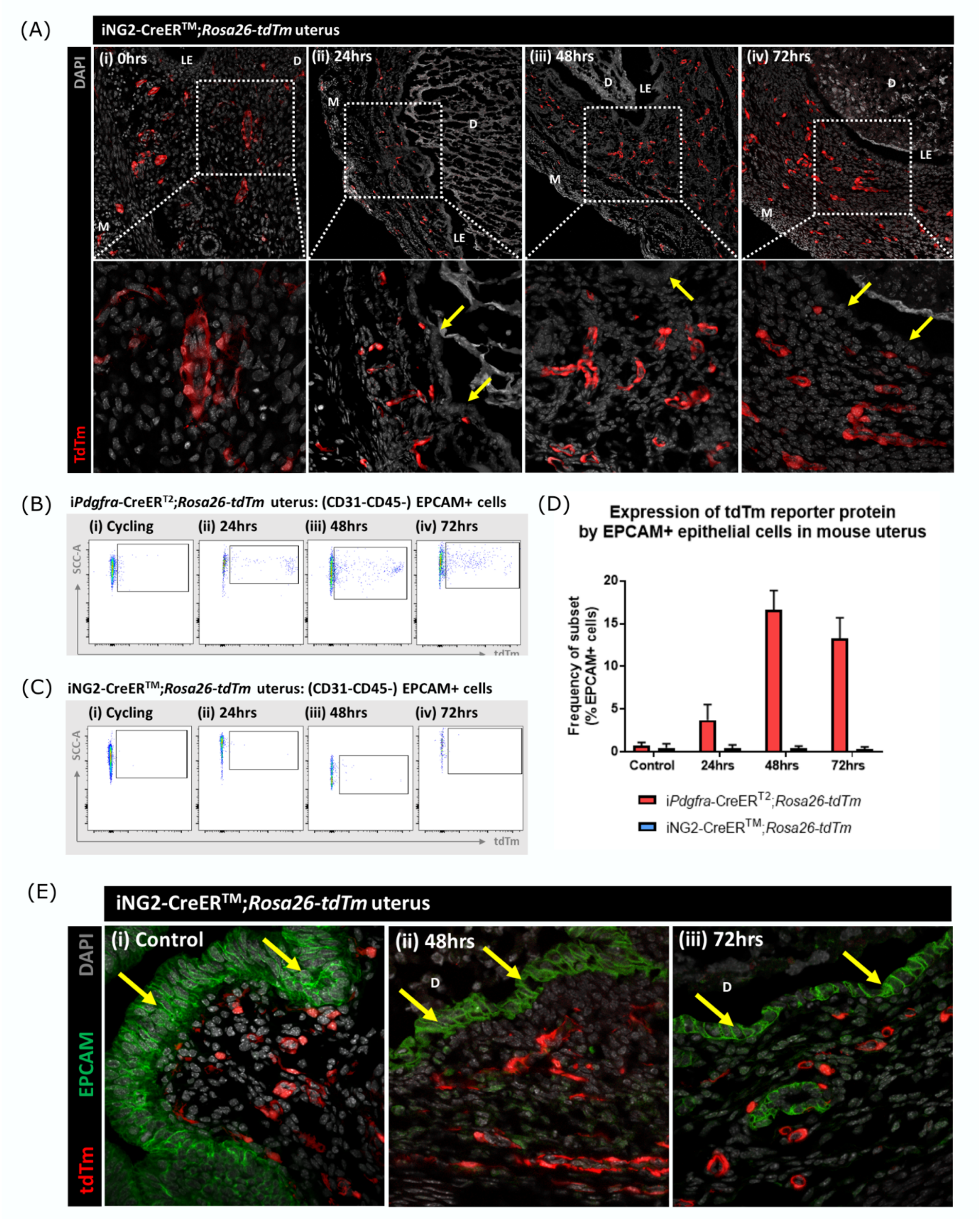


**Supplementary Figure 4. (A)** Analysis of tdTm reporter protein expression in iNG2-Cre$^{ERTM}$;*Rosa26-tdTm* uterine tissues 0, 24, 48 and 72h following progesterone withdrawal. Note tdTm+ cells were located throughout the stromal tissue but were never detected within the renewed luminal epithelium (yellow arrows) at any time point **(B)** Flow quantification of tdTm reporter expression by EPCAM+ cells in i*Pdgfra*-Cre$^{ERT2}$;*Rosa26-tdTm* uterine tissues 24, 48 and 72h following progesterone withdrawal. **(C)** Flow quantification of tdTm reporter expression by EPCAM+ cells in iNG2-Cre$^{ERTM}$;*Rosa26-tdTm* uterine tissues 24, 48 and 72h following progesterone withdrawal. **(D)** Bar plot: quantification of flow data analysing EPCAM+ cells that express tdTm in i*Pdgfra*-Cre$^{ERT2}$;*Rosa26-tdTm* or iNG2-Cre$^{ERTM}$;*Rosa26-tdTm* uterine tissues 24-72h following progesterone withdrawal. Note increase in the % of dpouble positive cells between 24 and 48h in the i*Pdgfra*-Cre$^{ERT2}$;*Rosa26-tdTm* but no evidence of co-staining with cells recovered from the iNG2 line; (**E)** Analysis of tdTm reporter and epithelial marker EPCAM expression in iNG2-Cre$^{ERTM}$;*Rosa26-tdTm* uterine tissues 48 and 72h following progesterone withdrawal. **(i-iii)** No co-localisation of tdTm and EPCAM was detected in steady state control tissues or 48/72hr tissues, tdTm+ cells were located throughout the stroma and EPCAM+ cells were located in the luminal and glandular epithelium (yellow arrows).
